# TRPM7 is critical for short-term synaptic depression by regulating synaptic vesicle endocytosis

**DOI:** 10.1101/2021.01.25.428048

**Authors:** Zhong-Jiao Jiang, Wenping Li, Li-Hua Yao, Brian S. Grewe, Andrea McGinley, Kelly Varga, Simon Alford, Liang-Wei Gong

## Abstract

TRPM7 contributes to a variety of physiological and pathological processes in many tissues and cells. With a widespread distribution in the nervous system, TRPM7 is involved in animal behaviors and neuronal death induced by ischemia. However, the physiological role of TRPM7 in CNS neuron remains unclear. Here, we identify endocytic defects in neuroendocrine cells and neurons from TRPM7 knockout (KO) mice, indicating a role of TRPM7 in synaptic vesicle endocytosis. Our experiments further pinpoint the importance of TRPM7 as an ion channel in synaptic vesicle endocytosis. Ca^2+^ imaging detects a defect in presynaptic Ca^2+^ dynamics in TRPM7 KO neuron, suggesting an importance of Ca^2+^ influx via TRPM7 in synaptic vesicle endocytosis. Moreover, the short-term depression is enhanced in both excitatory and inhibitory synaptic transmission from TRPM7 KO mice. Taking together, our data suggests that Ca^2+^ influx via TRPM7 may be critical for short-term plasticity of synaptic strength by regulating synaptic vesicle endocytosis in neurons.

## Introduction

Transient receptor potential melastatin 7 (TRPM7), an ubiquitously expressed member of transient receptor potential (TRP) superfamily (Nadler et al., 2001; Ramsey et al., 2006; Venkatachalam and Montell, 2007), is a cation channel fused with an unique alpha-kinase domain on its C terminus (Nadler et al., 2001). TRPM7 is able to transport divalent cations, particularly Mg^2+^ and Ca^2+^ (Monteilh-Zoller et al., 2003). TRPM7 is involved in physiological and pathological processes, including embryonic development, organogenesis, and organism survival (Jin et al., 2008; Ryazanova et al., 2010). The cellular functions of TRPM7 have been extensively studied in non-neuronal cells (Abumaria et al., 2018). The kinase domain of TRPM7 is believed to be important for cell growth and proliferation through either eEF2-kinase activation (Perraud et al., 2011) or chromatin remodeling (Krapivinsky et al., 2014). On the other hand, it has been suggested that the ion channel region of TRPM7 may be critical for cell survival by regulating Mg^2+^ homeostasis within cells, such as DT40 cell lines and embryonic stem cells (Ryazanova et al., 2010; Schmitz et al., 2003).

TRPM7 is widely distributed in the nervous system (Grube et al., 2003; Krapivinsky et al., 2006), and it may regulate animal behaviors, such as learning and memory in mouse and rat (Liu et al., 2018) and escape response in zebrafish (Low et al., 2011). A TRPM7 mutation with a defective ion channel property has been identified in human patients with Guamanian Amyotrophic Lateral Sclerosis and Parkinson’s disease (Hermosura et al., 2005), indicating a link between TRPM7 and neurodegenerative diseases. For the cellular physiology of neurons, TRPM7’s kinase domain is reported to be important for normal synaptic density and long-term synaptic plasticity in excitatory neurons from the central system (CNS) (Liu et al., 2018). The ion channel region of TRPM7 may regulate acetylcholine release in sympathetic neurons from the peripheral nervous system (PNS) (Krapivinsky et al., 2006; Montell, 2006), while the role of TRPM7 as an ion channel in CNS neurons has been focused on pathological neuronal death. TRPM7 may mediate neuronal Ca^2+^ overload, resulting in neuronal death under prolonged oxygen/glucose deprivation in neuronal culture (Aarts et al., 2003). Meanwhile, knocking down TRPM7 prevents neuronal death induced by either anoxia (Aarts et al., 2003) or ischemia (Sun et al., 2009). Furthermore, pharmacological blockage of TRPM7 reduces brain damage during neonatal hypoxic injury (Chen et al., 2015). TRPM7 as an ion channel is therefore considered as a potential molecular target for drug design to prevent neuronal death (Bae and Sun, 2013). Yet, the physiological role of the TRPM7’s ion channel region in CNS neurons remains unexplored.

A critical function of TRPM7 in membrane trafficking has been highlighted from non-neuronal cells (Abiria et al., 2017). Along this line, TRPM7 has been implicated in presynaptic vesicle recycling in both neuroendocrine cells (Brauchi et al., 2008) and PNS neurons (Krapivinsky et al., 2006). In PNS sympathetic neurons (Krapivinsky et al., 2006), it is reported that TRPM7 may regulate release of positively charged acetylcholine during vesicle fusion through a proposed ion compensation mechanism (Krapivinsky et al., 2006; Montell, 2006). However, it remains largely unknown what the physiological function of presynaptic TRPM7 is in CNS neurons. In the present study, by using a combination of biophysical, molecular biology, electrophysiological and live-cell imaging methods, the present study has identified that Ca^2+^ influx via TRPM7 may be critical for short-term changes of synaptic strength by regulating synaptic vesicle endocytosis in CNS neurons.

## Results

### Endocytic kinetics is slower in TRPM7 KO chromaffin cells

To understand potential roles of TRPM7 in exocytosis and endocytosis, TRPM7 knockout (KO) chromaffin cells were harvested from conditional KO newborn pups containing both Trpm7^fl/fl^ (Jin et al., 2008) and TH-Cre (a Cre specific to catecholaminergic cells (Savitt et al., 2005)) genes. The depletion of TRPM7 expression in medulla of adrenal glands was further confirmed by RT-PCR (Supplemental fig. 1A). Single vesicle endocytosis was monitored in neuroendocrine chromaffin cells using cell-attached capacitance recordings with a milli-second time resolution, and representative endocytic events from wildtype (WT) or TRPM7 KO cells were shown in Fig. 1A. Data from cell-attached capacitance recordings (Fig. 1A) showed that, while the number of endocytic events (Fig. 1B), the capacitance size of endocytic vesicles (Fig. 1C) and the fission-pore conductance (Gp) (Fig. 1D) were indistinguishable between WT and KO cells, the fission-pore duration (Fig. 1A & E) was significantly increased in TRPM7 KO cells, indicating a critical role of TRPM7 in regulating endocytic kinetics.

**Fig 1.**
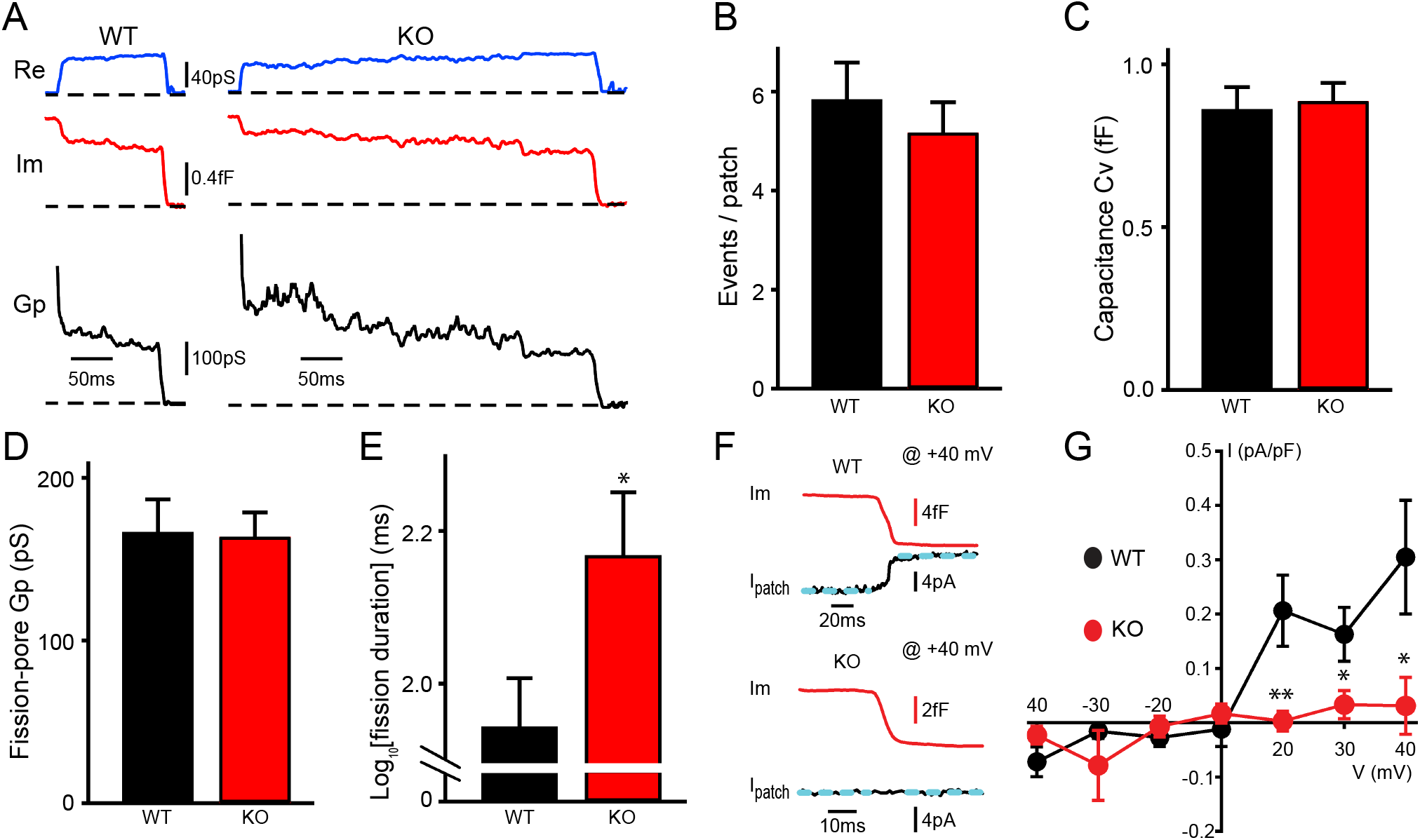
TRPM7 is important for endocytic fission during endocytosis in chromaffin cells. (**A**) Representative endocytic events, as membrane conductance (Re), membrane capacitance (Im) and the fission-pore conductance (Gp), in WT and TRPM7 KO cells. (**B**-**E**), The number of endocytic events (WT: n = 83 cells; KO: n = 110 cells) (B), the capacitance Cv of endocytic vesicles (C), and the fission-pore Gp (D) were statistically comparable between WT and KO cells; the log transformed fission-pore duration (E) was significantly increased in KO cells (WT: n= 42 events; KO: n= 42 events). (**F**) Representative membrane capacitance (Im) and patch membrane current (I_patch_) traces at +40 mV in WT and KO cells. Dashed lines in cyan represent linear fitting lines before or after capacitance drop induced by endocytosis. (**G**) The current-voltage relationship indicating that the endocytosis-associated current is reduced at positive membrane potentials in TRPM7 KO cells. n is 12, 9, 13, 12, 13, 10 and 11 for −40 mV, −30 mV, −20 mV, 0 mV, +20 mV, +30 mV and +40 mV in WT cells, and 12, 7, 8, 12, 11, 13 and 11 for −40 mV, −30 mV, −20 mV, 0 mV, +20 mV, +30 mV and +40 mV in KO cells, respectively. * p < 0.05 and * * p < 0.01, unpaired two-tailed student’s t-test.

Since localization of TRPM7 to synaptic vesicles has been indicated from previous studies (Brauchi et al., 2008; Krapivinsky et al., 2006), and to examine any potential TRPM7 openings during endocytosis, we performed double (whole-cell/cell-attached) patch recordings, in which the cell-attached patch pipette was utilized to monitor both endocytic events and the ionic current across the patch membrane. With the cell constantly clamped at - 60 mV by the whole-cell patch pipette, the patch of membrane captured by the cell-attached pipette was held at different potentials by varying the voltage in the cell-attached pipette. Fig. 1F showed at +40 mV an ionic current associated with endocytosis in WT cells but not in KO cells. The I-V relationships of the endocytosis-associated ionic current in WT cells (Fig. 1G) seemed to display the outwardly rectifying characteristics of whole-cell TRPM7 currents as reported in the literature (Nadler et al., 2001). Quantifications showed ionic currents at positive membrane potentials were substantially reduced in KO cells as compared to WT cells (Fig. 1G), suggesting TRPM7 openings during endocytosis. Interestingly, there was no such an ionic current associated with single vesicle exocytosis in WT cells (Supplemental fig. 2A), suggesting openings of TRPM7 channels specifically coincide with endocytosis but not exocytosis. Consistently, the biophysical calculation identified an extra conductance associated with endocytosis rather than exocytosis ((Supplemental fig. 2B-C).

**Fig 2.**
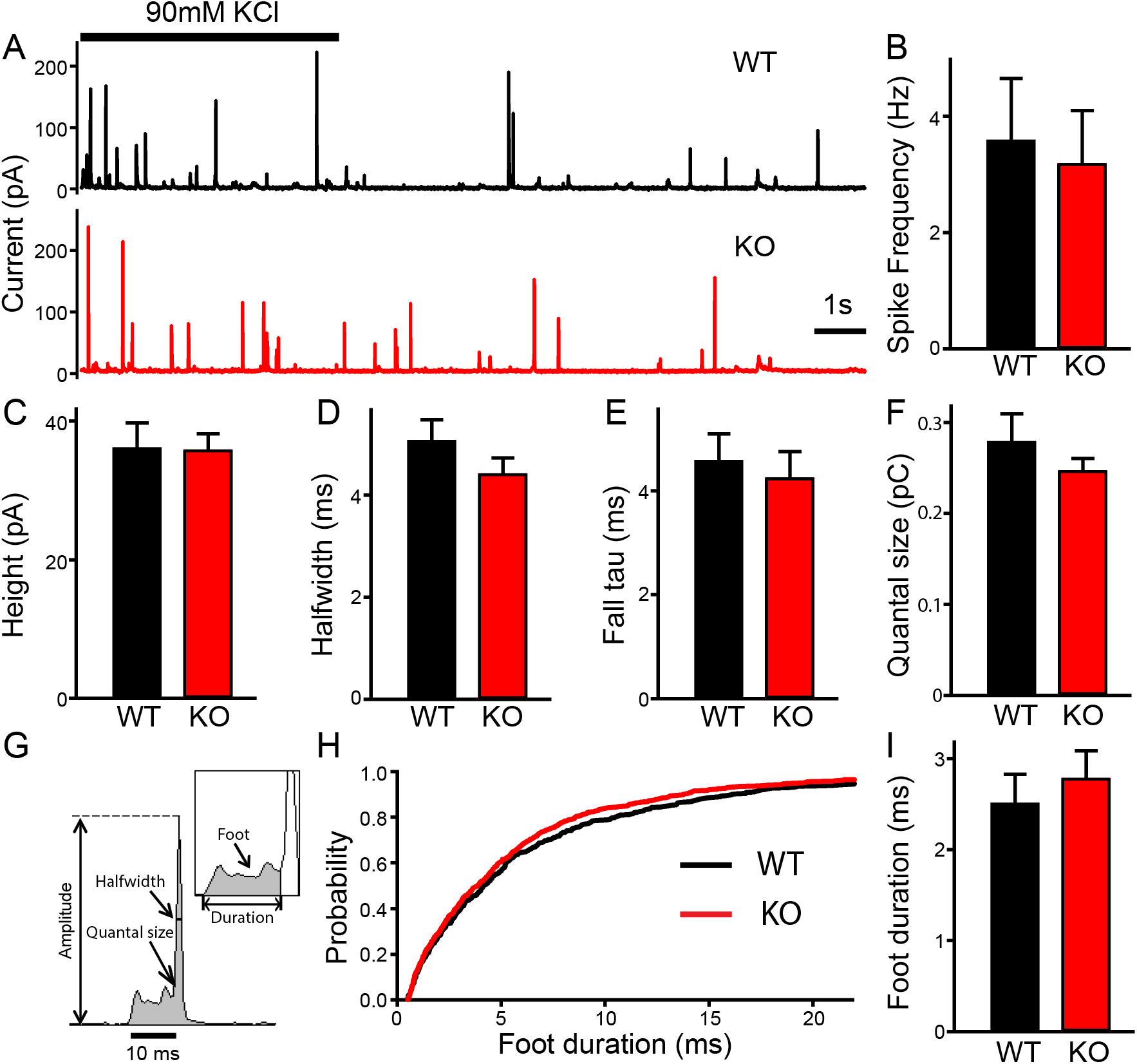
No change in LDCVs exocytosis in TRPM7 KO chromaffin cells detected by carbon fiber amperometry. (**A**) Amperometrical detection of catecholamine release from WT (upper) and KO (lower) chromaffin cells stimulated with 90 mM KCl for 5s. Each amperometrical spike represents release from an individual LDCV. (**B-F**) No differences between WT (n = 28) and TRPM7 KO (n = 31) cells in a variety of amperometric parameters, such as spike frequency (B), height (C), halfwidth (D), the exponential time constant of falling phase (E), and quantal size (F). (**G**). Diagram of the parameters analyzed in amperometric spikes. (**H-I**) Analysis of foot duration. Normalized cumulative distributions of all events in all cells (WT: n = 482; KO: n = 642) (H) and average of median values obtained from individual WT (n = 28) and KO (n = 31) cells (I).

### Exocytosis remains unaltered in TRPM7 KO chromaffin cells

To determine whether TRPM7 is involved in exocytosis, catecholamine release from single large dense-core vesicles (LDCVs) was analyzed using carbon fiber amperometry. Secretion induced by 90 mM KCl application for 5s was recorded by a carbon fiber electrode placed in a direct contact with cells (Fig. 2A). The frequency of amperometrical spikes within the first 15 s of stimulation was indistinguishable between WT and KO cells (Fig. 2B). Furthermore, no noticeable differences were observed between WT and KO cells for a variety of spike parameters such as peak amplitude (Fig. 2C), halfwidth (Fig. 2D), the time constant of falling phase (Fig. 2E) and quantal size (Fig. 2F). Amperometry can also provide detailed millisecond information about the initial release through a transient fusion pore (Chow et al., 1992), the foot signal, from the subsequent spike of catecholamine release (Fig. 2G). Our data showed no difference in foot duration between WT and TRPM7 KO cells (Fig. 2H-I). Therefore, our amperometrical results indicate TRPM7 may be dispensable for catecholamine release from LDCVs, which is consistent with a previous study showing that TRPM7 knockdown or expression of a dominant negative TRPM7 mutant does not affect LDCVs secretion in PC12 cells (Brauchi et al., 2008).

### Synaptic vesicle endocytosis is impaired in neurons from TRPM7 KO animals

To examine potential roles of TRPM7 in synaptic transmission, we cultured TRPM7 KO neurons from conditional KO newborn pups containing both Trpm7^fl/fl^ (Jin et al., 2008) and Nestin-Cre (a brain specific Cre (Tronche et al., 1999)) genes, and the depletion of TRPM7 in cortical neurons was confirmed by RT-PCR (Supplemental fig. 1B). Next, to directly examine synaptic vesicle endocytosis, we performed live-cell imaging of neurons with synaptophysin-pHluorin (sypHy) (Fernandez-Alfonso and Ryan, 2004; Granseth et al., 2006; Soykan et al., 2017; Voglmaier et al., 2006) expression driven by the human synapsin promoter (Nathanson et al., 2009; Yaguchi et al., 2013). There was an increase in the time constant of decay in sypHy fluorescent signals in TRPM7 KO neurons (Fig. 3A & B), indicative of an endocytic defect of synaptic vesicles. A similar endocytic defect was observed in neurons expressing vGlut1-pHluorin, which revealed a longer time constant for fluorescence decay in KO neurons as well (Fig. 3C-D). This endocytic defect cannot be the consequence from any changes in vesicle reacidification, since the reacidification rate of newly formed endocytic vesicles was comparable between WT and KO neurons measured with sypHy (Fig. 3E-F). Collectively, our data indicates that TRPM7 may be important for synaptic vesicle endocytosis in neurons.

**Fig 3.**
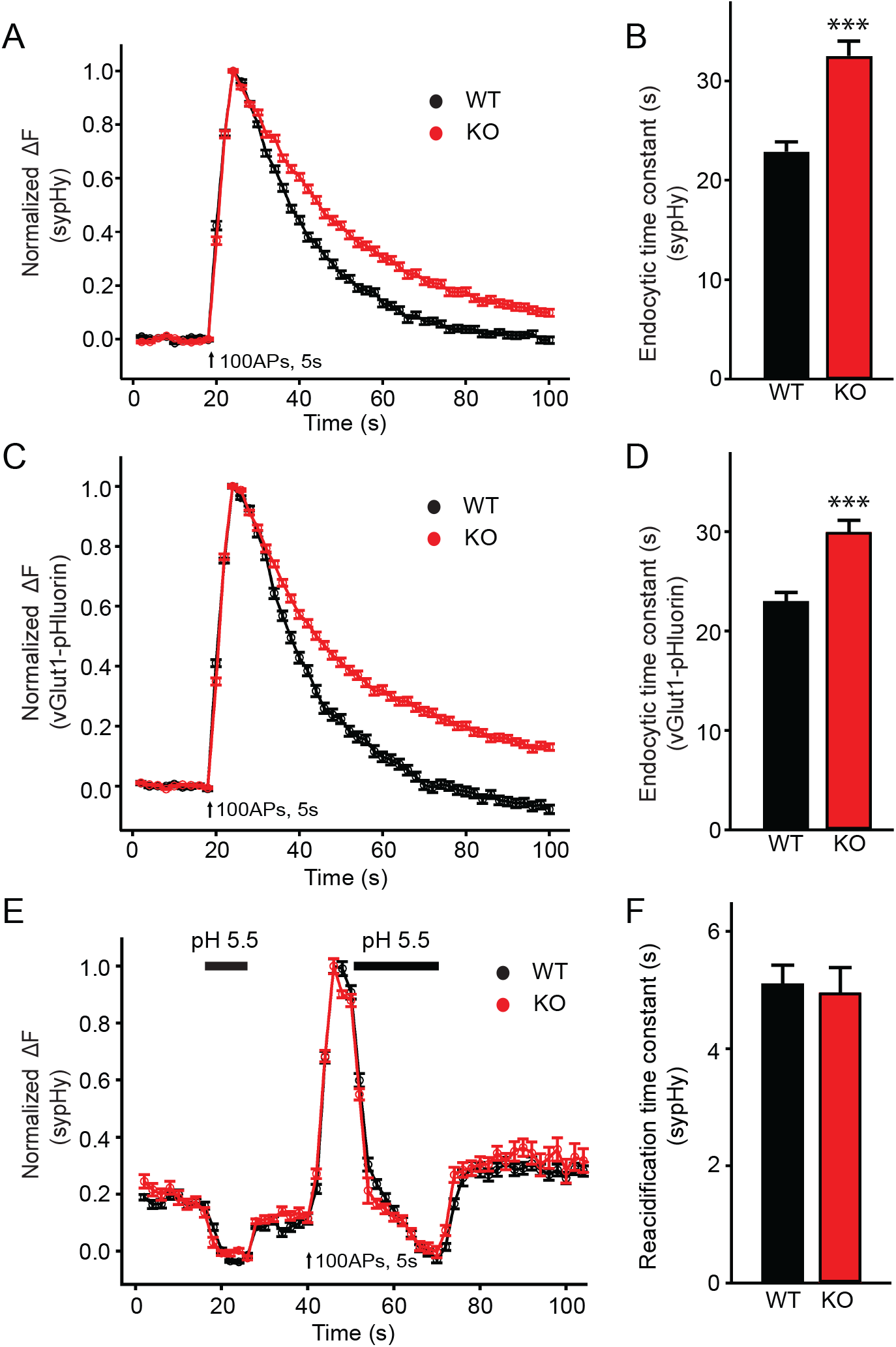
Synaptic vesicle endocytosis is impaired in TRPM7 KO neurons using pHluorin-based optical imaging assays. (**A**) Normalized fluorescence changes of sypHy signals in WT (n = 139) and KO (n= 127) neurons at 2 mM [Ca^2+^]_e_. (**B**) Bar graph comparing the endocytic time constant from sypHy experiments in a. (**C**) Normalized fluorescence changes of vGlut1-pHluorin signals in WT (n = 72) and KO (n= 80) neurons at 2 mM [Ca^2+^]_e_. (**D**) Bar graph comparing the endocytic time constant from vGlut1-pHluorin experiments in c. (**E**) Normalized sypHy fluorescence traces (WT: n = 61; KO: n = 45) with two periods of perfusion with pH 5.5 MES buffer as indicated in the figure (black bar). Newly endocytosed vesicles “trapped” during acidic buffer perfusion starting 5 s after 100 APs at 20 Hz. (**F**) Bar graph for the reacidification rate of endocytic vesicles, which was obtained by exponential fittings of fluorescence decay, indicates no change in reacidification in KO neurons. n is the total number of boutons pooled from 3 animals. * * * p < 0.001, unpaired two-tailed student’s t-test.

Conversely, paired whole-cell recordings in pyramidal neurons of cortical culture revealed no change in the amplitude of IPSCs evoked by single presynaptic stimulation with 1-ms depolarization to +30 mV in KO neurons (Supplemental fig. 3A-B). There was also no obvious change in the paired pulse ratio (Supplemental fig. 3C), indicating no alteration in release probability in KO neurons. These results suggest that TRPM7 may not be critical for exocytosis of synaptic vesicles in inhibitory synapses with electrically neutral GABA as the predominant neurotransmitter, although it is aware that a previous study implies that TRPM7 may have a post-fusion role by supplying counterions during release of positively charged acetylcholine in PNS sympathetic neurons (Krapivinsky et al., 2006; Montell, 2006).

### WT TRPM7 (TRPM7^WT^), but not a non-conducting TRPM7 mutant (TRPM7^LCF^), rescues endocytic defects in TRPM7 KO chromaffin cells and neurons

To further explore whether the ion channel region of TRPM7 is relevant to TRPM7-dependent synaptic vesicle endocytosis, we generated a non-conducting TRPM7 mutant with a loss of ion channel functions (TRPM7^LCF^), by mutating highly conserved amino acids located in the sixth putative transmembrane domain (amino acids 1090–1092, NLL mutated to FAP) (Krapivinsky et al., 2006). As reported previously (Krapivinsky et al., 2006), when compared to TRPM7^WT^, there was an almost complete suppression of TRPM7^LCF^ currents in HEK293 cells with lentiviral infection (Fig. 4A-B and supplemental fig. 4). Using cell-attached recordings, we examined kinetics of single endocytic events in KO chromaffin cells expressing either TRPM7^WT^ or TRPM7^LCF^ (Fig. 4C), with similar TRPM7^WT^ or TRPM7^LCF^ expression levels as confirmed by immunostaining (Supplemental fig. 5). While the number of endocytic events (Fig. 4D), the capacitance size of endocytic vesicles (Fig. 4E), and the fission-pore Gp (Fig. 4F) were comparable between these two groups; the fission-pore duration was reduced in KO cells expressing TRPM7^WT^ as compared to TRPM7^LCF^ (Fig. 4G). Consistently, the time constant of sypHy signal decay was shorter in KO neurons expressing TRPM7^WT^ than TRPM7^LCF^ from live-cell imaging experiments (Fig. 4H-I). These findings thus conclude that TRPM7’s ion channel region may be important for synaptic vesicle endocytosis in neurons.

**Fig 4.**
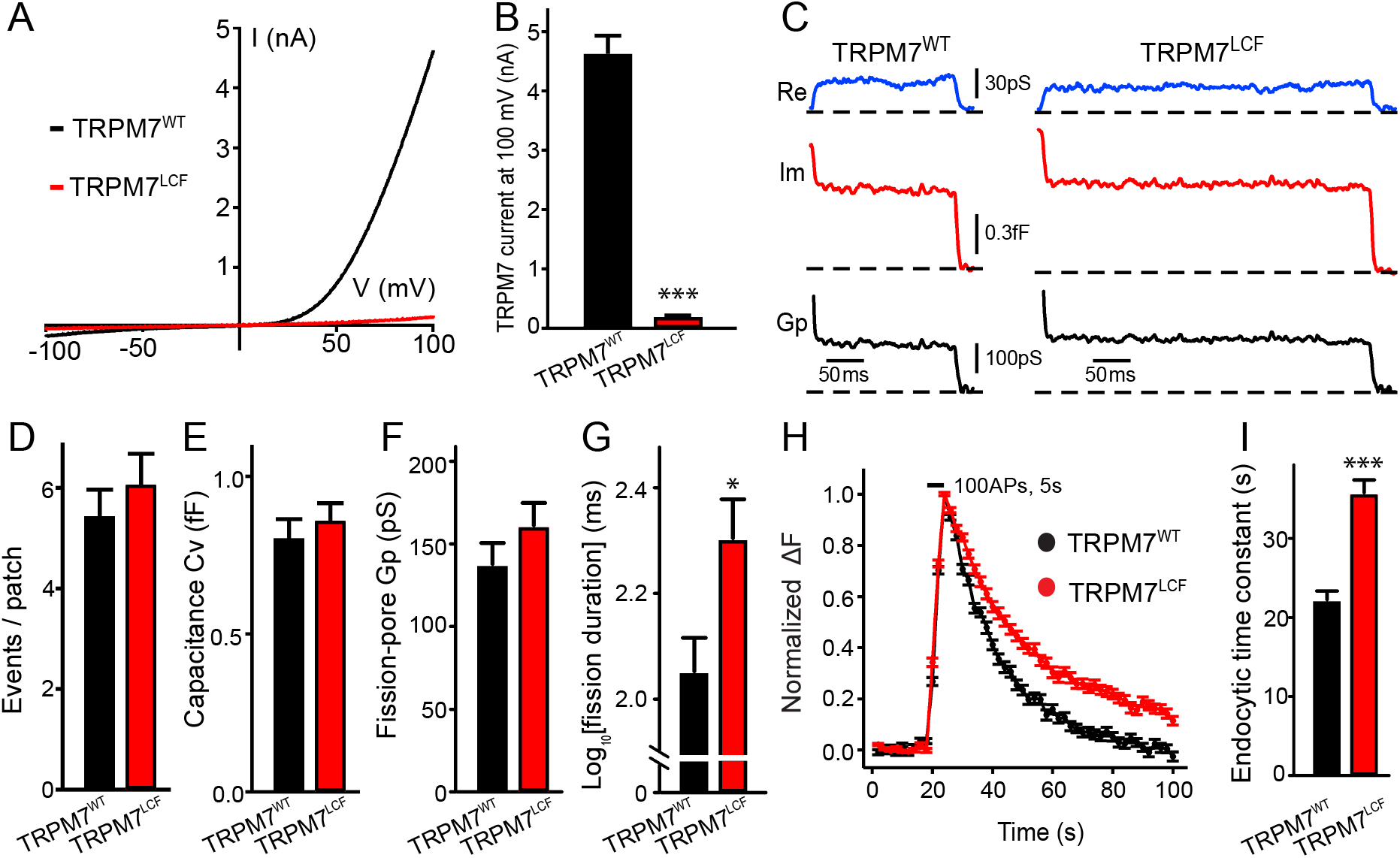
TRPM7 as an ion channel is critical for endocytosis in both chromaffin cells and neurons. (**A**) Averaged whole-cell TRPM7 currents, recorded from HEK293 cells transduced with TRPM7^WT^ or TRPM7^LCF^, in response to voltage ramps from −100 mV to +100 mV. (**B**) Quantifications showing TRPM7^LCF^ currents (n = 14 cells) at + 100 mV were almost completely blocked as compared to TRPM7^WT^ currents (n = 14 cells). (**C**) Representative endocytic events, as membrane conductance (Re), capacitance (Im) and the fission-pore Gp traces in KO cells expressing either TRPM7^WT^ (left) or TRPM7^LCF^ (right). (**D**-**G**) The number of endocytic events (TRPM7^WT^: n = 109 cells; TRPM7^LCF^: n = 86 cells) (D), the capacitance Cv of endocytic vesicles (E), and the fission-pore Gp (F) were indistinguishable between these 2 groups; the log transformed fission-pore duration (G) was significantly increased in TRPM7^LCF^-expressing KO cells (TRPM7^WT^: n = 44 events; TRPM7^LCF^: n = 53 events). (**H**-**I**) Normalized fluorescence changes of sypHy signals (H) and bar graph (I), comparing the endocytic time constant from sypHy experiments in KO neurons expressing TRPM7^WT^ (n = 69) or TRPM7^LCF^ (n = 87) neurons, indicates that endocytic kinetics is slower in TRPM7^LCF^-expressing KO neurons. n is the total number of boutons pooled from 6 animals. * p < 0.05 and * * * p < 0.001, unpaired two-tailed student’s t-test.

**Fig 5.**
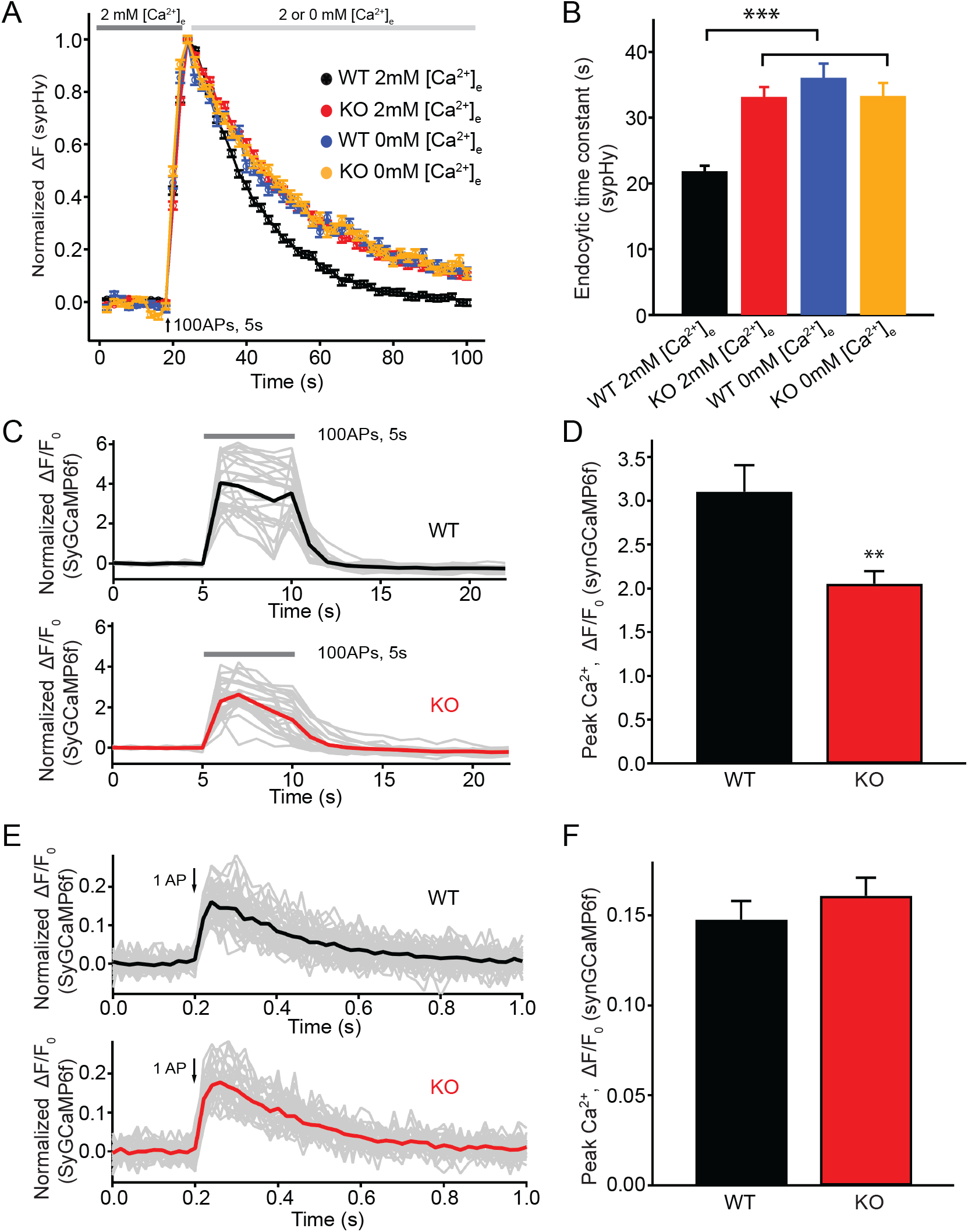
TRPM7 is important for presynaptic Ca^2+^ increase upon a train of stimulations. (A) Normalized fluorescence changes of sypHy signal in WT (2 mM: n = 131; 0 mM: n = 68) and TRPM7 KO (2 mM: n= 116; 0 mM: n = 67) neurons at 2 and 0 mM [Ca^2+^]_e_ after the cessation of stimuli. (**B**) Bar graph comparing the endocytic time constant from sypHy experiments in a. (**C**) Sample traces showing normalized changes in SyGCaMP6f fluorescence stimulated with 100 APs at 20 Hz from an individual coverslip of WT (top) (25 ROIs) or TRPM7 KO (bottom) (25 ROIs) neurons at 2 mM [Ca^2+^]_e_. Gray lines are changes for individual ROIs with the black or red line as the averaged response. (**D**) Bar graph comparing peak values of Ca^2+^ transients from SyGCaMP6f experiments in c displays a significant decrease of Ca^2+^ signals in TRPM7 KO neurons (WT: n=11; KO: n=10). (**E**) Sample traces showing normalized changes in SyGCaMP6f fluorescence induced by single AP from an individual coverslip of WT (top) (36 ROIs) or TRPM7 KO (36 ROIs) neurons at 2 mM [Ca^2+^]_e_. Gray lines are changes of individual ROIs with black or red line as the averaged response. (**F**) Bar graph comparing peak values of Ca^2+^ transients as shown in e revealed no change in Ca^2+^ signals induced by a single AP (WT: n=22; KO: n=22) (p > 0.01). * * p < 0.01, unpaired two-tailed student’s t-test; * * * p < 0.001, Newman of one-way ANOVA in b.

### The presynaptic Ca^2+^ increase upon a train of stimulations is reduced in TRPM7 KO neurons

Our sypHy experiments showed that there was no difference in the endocytic time constant of fluorescent signal decay between WT and KO neurons perfused with 0 mM [Ca^2+^]_e_ right after the cessation of stimulations (Fig. 5A-B), suggesting a role of TRPM7 in Ca^2+^- dependent rather than Ca^2+^-independent synaptic vesicle endocytosis. Therefore, it is speculated that Ca^2+^ influx via TRPM7 may be critical for synaptic vesicle endocytosis. To test this idea, we compared presynaptic Ca^2+^ signaling between WT and TRPM7 KO neurons by using Synaptophysin-GCaMP6f (SyGCaMP6f) fusion protein as a presynaptic Ca^2+^ reporter (de Juan-Sanz et al., 2017). While neurons responded to electrical stimuli with a robust increase in SyGCaMP6f fluorescence (Fig. 5C-D), the increase in the SyGCaMP6f signal was substantially reduced in response to a stimulation of 100 APs at 20 Hz in TRPM7 KO neurons (Fig. 5C-D). The reduction in Ca^2+^ rise in response to stimulation train is unlikely due to any changes in Ca^2+^ influx via voltage gated Ca^2+^ channels (VGCCs), since the peak of SyGCaMP6f signals induced by single AP was indistinguishable between WT and KO neurons (Fig. 5E-F) (Brockhaus et al., 2019; Kim and Ryan, 2013).

### Short-term depression of synaptic transmission is enhanced in TRPM7 KO neurons

Our data presented so far indicates that Ca^2+^ influx via TRPM7 may be critical for synaptic vesicle endocytosis. Since synaptic vesicle endocytosis is an essential presynaptic factor for short-term synaptic plasticity (Regehr, 2012; Zucker and Regehr, 2002), we next examined whether short-term synaptic depression is altered in TRPM7 KO neurons.

For inhibitory synaptic transmission, evoked IPSCs in pyramidal neurons were recorded by stimulating presynaptic inhibitory neurons with 250 stimuli at 10 Hz through a presynaptic whole-cell pipette (Fig. 6A). With the amplitude of evoked IPSCs normalized to the first response, KO neurons displayed a more profound short-term synaptic depression (Fig. 6A-B) and a 56% reduction of the steady-state IPSCs (insert in Fig. 6B), suggesting an important role of TRPM7 in the short-term synaptic depression of inhibitory synaptic transmission.

**Fig 6.**
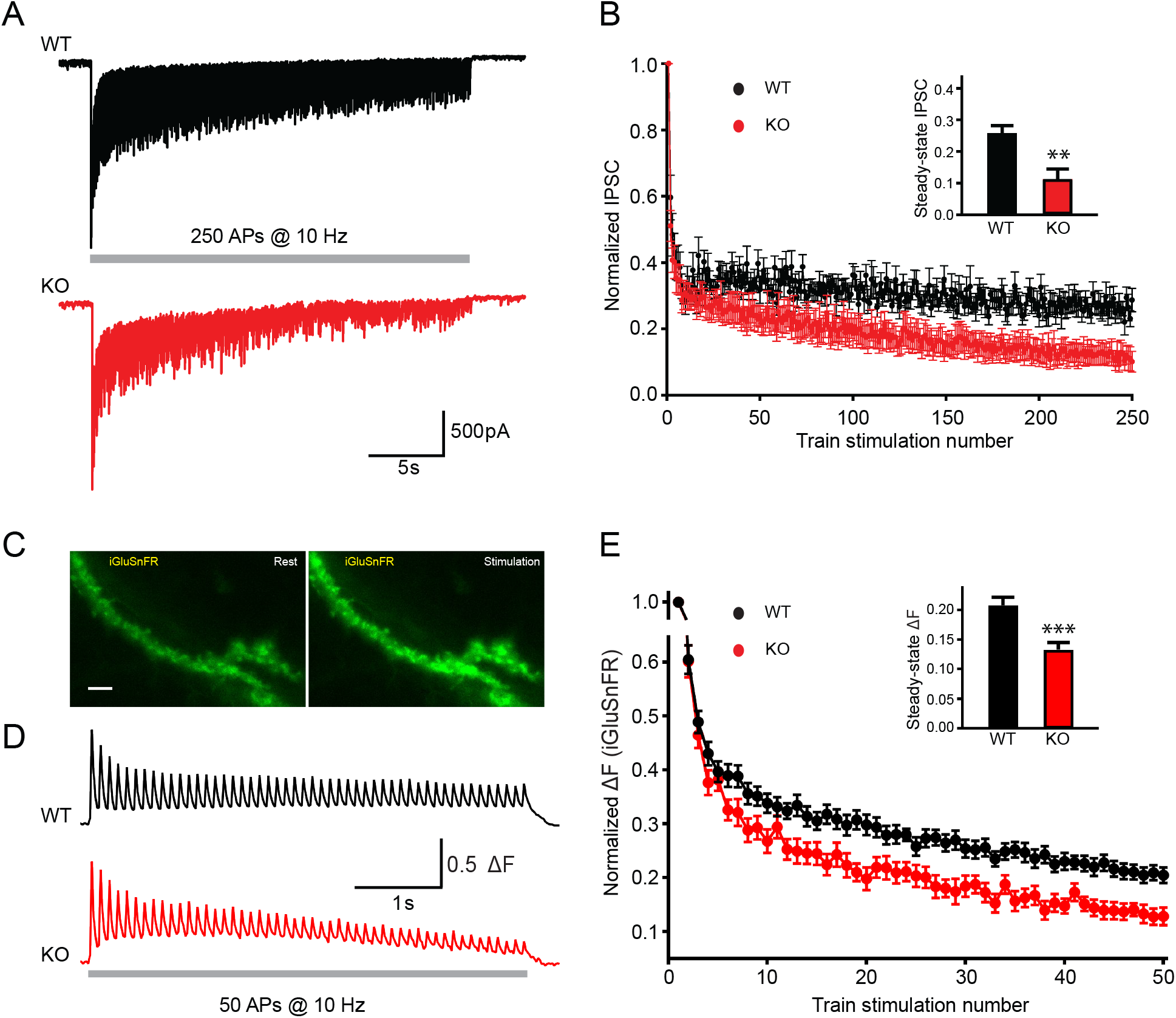
TRPM7 is critical for presynaptic short-term depression in both inhibitory and excitatory synaptic transmissions. (**A**-**B**) Representative traces of IPSCs evoked by 250 stimulations at 10 Hz in WT (top in a) and TRPM7 KO (bottom in a) neurons (A) and graph of averaged normalized IPSCs changes (B). Bar graph of the steady-state IPSCs calculated as the average of last 10 evoked responses in WT and TRPM7 KO neurons are shown in insert in b. Number of recordings is 10 for WT and 8 for KO. (**C**) Representative images showing fluorescence levels of the same dendritic arbor expression iGluSnFR at rest (left) or upon stimulations (right). Scale bar, 5 µm. (**D-E**) Representative traces, from WT (top in d) and TRPM7 KO (bottom in d) neurons (D), and normalized graphs (E) of changes in iGluSnFR fluorescence (ΔF/ΔFmax) evoked by 50 stimulations at 10 Hz. Bar graph of the steady-state of iGluSnFR ΔF/ΔFmax calculated as the average of last 5 evoked responses in WT and TRPM7 KO neurons are shown in insert in e. Number of coverslips is 21 for WT and 20 for KO. * * p < 0.01, and * * * p < 0.001, unpaired two-tailed student’s t-test.

For excitatory synaptic transmission, high-speed optical imaging of evoked glutamate release using iGluSnFR (Marvin et al., 2018) was utilized to directly monitor the presynaptic short-term plasticity (Fig. 6C). Neurons on coverslips with iGluSnFR expression were challenged with 50 stimuli at 10 Hz to induce short-term depression of evoked glutamate release, fluorescent healthy dendrite arbors were focused and imaged at 50 Hz (Fig. 6C-D). Similar to that observed in the synaptic plasticity measured with evoked IPSCs (Fig. 6A-B), normalized changes of the iGluSnFR fluorescence during the train also revealed an enhanced short-term depression in TRPM7 KO neurons (Fig. 6D-E). The normalized steady-state iGluSnFR signals at the end of the train was reduced by 36% in TRPM7 KO neurons (p < 0.001) (insert in Fig. 6E), implying a significance of TRPM7 in short-term depression of excitatory synaptic transmission.

Taking together, the enhanced presynaptic short-term depression in both inhibitory and excitatory neurons from TRPM7 KO animals (inserts in Fig. 6B & E) corroborated each other. Given the well-established significance of synaptic vesicle endocytosis in short-term depression^33,34^, our data may propose that TRPM7 is critical for presynaptic short-term synaptic depression via its regulation in synaptic vesicle endocytosis.

## Discussion

### TRPM7 may serve as a Ca^2+^ influx pathway for synaptic vesicle endocytosis

Since VGCCs have been shown to be important for endocytosis (Midorikawa et al., 2014; Perissinotti et al., 2008; Xue et al., 2012), it is implied that Ca^2+^ for endocytosis at the periactive zone may be derived from diffusion of VGCCs-mediated Ca^2+^ influxes at active zones. On the other hand, exocytosis can be triggered, without activations of VGCCs, by black widow spider venom (Ceccarelli and Hurlbut, 1980; Henkel and Betz, 1995), caffeine (Zefirov et al., 2006), or veratridine (Kuromi and Kidokoro, 2002). Interestingly, extracellular Ca^2+^ is still necessary for synaptic vesicle endocytosis in these paradigms, suggesting distinct Ca^2+^ channels independent of VGCCs for endocytosis. Indeed, a La^3+^- sensitive Ca^2+^ channel is indicated to be required for synaptic vesicle endocytosis in *Drosophila* (Kuromi et al., 2004).

While it has been proposed that a Ca^2+^-permeable Flower channel may be necessary for synaptic vesicle endocytosis at the neuromuscular junction in *Drosophila* (Yao et al., 2009), the function of Flower as the Ca^2+^ channel in synaptic vesicle endocytosis of mammalian systems remains questionable, since the role of this protein in synaptic vesicle endocytosis was not confirmed at the calyx of Held (Xue et al., 2012). Additionally, a recent study showed that the protein itself but not the ion channel property of Flower is important for synaptic vesicle endocytosis elicited by moderate stimulations in rat hippocampal neurons (Yao et al., 2017). Furthermore, the time course of Ca^2+^ influx via Flower protein, in the order of 10-60 min (Yao et al., 2009), may be too slow for synaptic vesicle endocytosis.

Collectively, our data suggest that TRPM7 channels may serve a Ca^2+^ influx pathway for synaptic vesicle endocytosis in cortical neurons. Along this line, it is of note that Ca^2+^ influx via TRPM7 may be essential for endocytosis of toll-like receptor 4 in the non-neuronal microphage (Schappe et al., 2018). A recent study indicates that TRPC5, another TRP superfamily member, may serve as a Ca^2+^ influx pathway for synaptic vesicle endocytosis in hippocampal neurons (Schwarz et al., 2019). Since TRP channels are widely expressed in the brain (Kunert-Keil et al., 2006), it is speculated that members of the TRP superfamily may serve as general Ca^2+^ channels for synaptic vesicle endocytosis in different regions of the nervous system.

### Ca^2+^ influx via TRPM7 may regulate the kinetics rather than the number of single endocytic events

The rate of synaptic vesicle endocytosis is determined by the number of vesicles involved and the kinetics of single endocytic events. Our results, showing slower kinetics but no change in the number of single endocytic events in TRPM7 KO chromaffin cells, indicate that Ca^2+^ influx via TRPM7 may regulate the rate of synaptic vesicle endocytosis by regulating the kinetics of single endocytic events but not the number of vesicles involved. This result is in line with our previous study, which demonstrates an importance of extracellular Ca^2+^ for the kinetics but not the number of single vesicle endocytosis (Yao et al., 2012). Consistently, extracellular Ca^2+^ may modulate endocytic kinetics of single synaptic vesicles in neurons, although no analysis was performed on whether Ca^2+^ affects the number of synaptic vesicles involved (Leitz and Kavalali, 2011).

### TRPM7 may be a vesicular protein in presynaptic terminals

By simultaneously monitoring ionic current and membrane capacitance in neuroendocrine chromaffin cells, we have identified an endocytosis-associated ionic current, which is substantially reduced in TRPM7 KO cells (Fig. 1F-G). Since the endocytosis-associated ionic current drifts rather than displaying a pattern of single channel openings (Fig. 1F), we reason that the endocytosis-associated ionic current may be attributed to a loss of multiple small conductance ion channels on the vesicle when the endocytic vesicle detaches from the plasma membrane. Consistently, our data shows that the endocytosis-associated current drift is synchronized with capacitance drop, which reflects a separation of the endocytic vesicle from the plasma membrane. In fact, TRPM7 is a channel that can be activated by PI(4,5)P_2_ and inactivated upon PI(4,5)P_2_ hydrolysis (Runnels et al., 2002). Interestingly, PI(4,5)P_2_ is concentrated on the plasma membrane but absent from synaptic vesicles, due to PI(4,5)P_2_ hydrolysis by phosphatases such as synaptojanin 1 during uncoating after vesicle pinch-off, a late stage of endocytosis (Cremona and De Camilli, 2001). Therefore, the specific openings of TRPM7 during endocytosis may be attributed to the tight coupling between PI(4,5)P_2_ metabolism and synaptic vesicle recycling (Cremona and De Camilli, 2001).

Consistent with our data, previous studies have indicated TRPM7 as a vesicular protein in non-neuronal cells (Abiria et al., 2017), neuroendocrine cells (Brauchi et al., 2008), and neurons (Krapivinsky et al., 2006). Immunostaining in PC12 cells, a cell line of neuroendocrine chromaffin cells, has found that TRPM7 is localized to small synaptic-like vesicles (Brauchi et al., 2008). A biochemical analysis of purified rat brain synaptosomes and synaptic vesicles supports this finding by revealing that TRPM7 is concentrated in those regions but absent from postsynaptic densities (Krapivinsky et al., 2006). Further, the same study suggests that TRPM7 may form a molecular complex with vesicular proteins such as synapsin 1 and synaptotagmin 1 (Krapivinsky et al., 2006). On the other hand, a systematic survey of synaptic vesicle proteins did not detect TRPM7 as a synaptic vesicle protein (Takamori et al., 2006). However, this survey has missed some well-known synaptic vesicle proteins, such as SVOP, the chloride channels CIC3 and CIC7, and ever the putative vesicular neurotransmitter transporters (VMATs and VACHT). Indeed, as claimed by the authors, their proteomics results in an overall coverage of ∼ 80% of all known vesicle membrane proteins (Takamori et al., 2006). Therefore, it is possible that the systematic survey of synaptic vesicle proteins may have failed to detect TRPM7 on synaptic vesicles.

### The physiological role of TRPM7 in CNS neurons

As an ion channel fused with a c-terminal alpha-kinase (Nadler et al., 2001), TRPM7’s ion channel region may be critical for synaptic vesicle endocytosis (the present study) while its kinase domain may be important for normal synaptic density (Liu et al., 2018) in CNS neurons under physiological conditions. It is reported that TRPM7 may be involved in learning and memory in mouse and rat, as knockdown or conditional knockout of TRPM7 impairs spatial learning (Liu et al., 2018). The authors indicate that the role of TRPM7 in learning and memory could reflect its importance in regulating synaptic density and long-term synaptic plasticity in excitatory neurons (Liu et al., 2018). However, this interpretation is conflicting with a previous study showing that TRPM7 knockdown does not change evoked synaptic response, an indicative of no change in synaptic density, and LTP (Sun et al., 2009). Interestingly, recent studies suggest that TRPC5, which regulates synaptic vesicle endocytosis through a mechanism (Schwarz et al., 2019) similar to TRPM7 as we have observed in the present study, is critical for both short-term synaptic plasticity in excitatory neurons (Schwarz et al., 2019) and learning and memory of animals (Broker-Lai et al., 2017). Therefore, the role of TRPM7 in short-term synaptic plasticity in excitatory neurons identified here may presumably account for its function in learning and memory of animals reported previously (Liu et al., 2018). Notably, physiological roles of TRPM7 in inhibitory neurons remain largely unknown, although TRPM7 is actually expressed in both excitatory and inhibitory neurons from single cell transcriptome (Huntley et al., 2020). Our study identifies that TRPM7 is also important for short-term synaptic plasticity in inhibitory neurons. Given the increasing significance of inhibitory neuronal circuits in learning and memory (Barron et al., 2017), the reported roles of TRPM7 in learning and memory (Liu et al., 2018) could, alternatively, be attributed to its modulations of inhibitory synaptic transmission.

### Implications

In summary, our study indicates that Ca^2+^ influx via TRPM7 is critical for synaptic vesicle endocytosis and consequently short-term synaptic depression, thus implying that TRPM7 may critical for filtering and gain control of synaptic transmission, as short-term synaptic plasticity is an important factor of synaptic computation (Abbott and Regehr, 2004). TRPM7 is a Mg^2+^-regulated ion channel and the Mg^2+^-dependent modulation can be relieved by membrane depolarizations (Ramsey et al., 2006), a disinhibition mechanism similar to NMDA receptors. By identifying a role of TRPM7 as an ion channel in synaptic vesicle endocytosis, it is reasonable to speculate that TRPM7 may serve as a molecule that couples neuronal activity to synaptic vesicle endocytosis.

Recent studies have revealed a tight link between neurodegenerative diseases, particularly Parkinson’s disease, and proteins that participate in synaptic vesicle endocytosis, such as auxilin (Song et al., 2017), synaptojanin 1 (Cao et al., 2017) and endophilin (Trempe et al., 2009). Our evidence for a role of TRPM7 in synaptic vesicle endocytosis suggests that the consequence on synaptic vesicle endocytosis attributed to TRPM7 dysfunctions could be a potential molecular mechanism for TRPM7 related neurodegenerative diseases, such as Guamanian Amyotrophic Lateral Sclerosis and Parkinson’s disease (Hermosura et al., 2005).

## Materials and Methods

### Generation of TRPM7 conditional KO mice and mouse breeding

All animal experimental studies were approved by the Animal Care and Use Committee of the University of Illinois at Chicago and conformed to the guidelines of the National Institutes of Health. All mating cages were exposed to a 12-h light/dark cycle with food and water provided *ad libitum*.

TRPM7^fl/fl^ (#018784, a kind gift of Dr. Clapham lab (Jin et al., 2008)), Nestin-Cre (#003771, a kind gift from Dr. Schütz lab (Tronche et al., 1999)) and TH-Cre (#008601, a kind gift from Dr. Dawson lab (Savitt et al., 2005)) mice were obtained from Jackson lab. TRPM7^fl/fl^ mice were crossed with TH-Cre or Nestin-Cre line to produce hemizygous TH-Cre or Nestin-Cre mice, which were heterozygous for the loxP targeted (floxed) *Trpm7* gene (TH-Cre/TRPM7^fl/+^ or Nestin-Cre/TRPM7^fl/+^), respectively. Male TH-Cre/TRPM7^fl/+^ mice were then backcrossed to female TRPM7^fl/fl^ mice to generate mice with TRPM7 KO catecholaminergic cells including chromaffin cells (TH-Cre/TRPM7^fl/fl^). Similarly, Nestin-Cre/TRPM7^fl/+^ mice were backcrossed to TRPM7^fl/fl^ mice to generate brain specific TRPM7 KO mice (Nestin-Cre/TRPM7^fl/fl^). Controls were the littermates carrying a single floxed allele of TRPM7 (TRPM7^fl/+^).

Cre transgenes were genotyped using tail biopsy DNA samples with PCR primers 5’-GCGGTCTGGCAGTAAAAACTATC-3’ and 5’-GTGAAACAGCATTGCTGTCACTT-3’ to detect a 100 bp transgenic fragment. The TRPM7 floxed and WT alleles were detected using the primer set: forward, 5′-TTTCTCCAATTAGCCCTGTAGA-3′; reverse, 5′-CTTGCCATTTTACCCAAATC-3′, the products were 300 bp for the floxed allele and 193 bp for the WT allele. RT-PCR of adrenal medulla and cerebral cortex mRNA were used to confirm the excision of loxP-flanked (floxed) sequences, indicated by a 454 bp cDNA fragment of wild-type TRPM7, and a 225 bp fragment from Cre recombinase-excised, as previously described (Jin et al., 2008).

### Cultures of chromaffin cells, neurons and HEK 293 cells

Chromaffin cells in culture, prepared from adrenal glands of newborn pups with either sex as previously described (Gong et al., 2005; Yao et al., 2013; Yao et al., 2012), were maintained at 37°C in 5% CO_2_ and used within 4 days for electrophysiology or immunostaining. Lentiviral infections with TRPM7^WT^ or TRPM7^LCF^ in TRPM7 KO cells were carried out at *days in vitro* (DIV) 0, and electrophysiological recordings or immunostaining staining was performed 72 to 96 hrs after infection.

Neuronal culture from cortex of newborn pups with either sex was prepared as described previously (Gong and De Camilli, 2008). Neurons in culture were maintained at 37°C in 5% CO_2_ and used at DIV 14-16 for electrophysiology. Neurons at DIV 5 were transfected with vGlut1-pHluorin, TRPM7^WT^-sypHy or TRPM7^LCF^-sypHy plasmids using lipofectamine LTX (Thermo Fisher) or infected with sypHy, SyGCaMP6f, or iGluSnFR lentivirus, and imaging experiments were carried out on neurons at DIV 16-20.

HEK293 cells were cultured in DMEM supplemented with 10% FBS, 0.1 mM MEM non-essential amino acids, 6 mM L-glutamine, 1 mM MEM sodium pyruvate, 1% pen-strep and 500 ug/ml Geneticin™ antibiotic. Cells, which typically reached 80% confluence every 3 days, were treated with trypsin-EDTA and passaged with a 1:10 ratio for the general maintenance. Cells at low density were seeded onto 24 well culture plates and infected with lentivirus carrying FLAG tagged TRPM7^WT^ or TRPM7^LCF^, and monoclonal cell lines were then generated by limiting dilution. After confirming the monoclonal cell lines expressing either TRPM7^WT^ or TRPM7^LCF^ by FLAG immunostaining, the monoclonal cell lines were maintained at 37 °C in a 5% CO_2_ incubator for 48-72 hours before being trypsinized and seeded to PDL coated coverslip for whole-cell patch clamping recordings.

### Cell-attached capacitance recordings and analyses of exocytic fusion-pore and endocytic fission-pore

Cell-attached capacitance recordings of membrane capacitance and conductance were performed as described previously (Yao et al., 2013; Yao et al., 2012). Fire polished pipettes had a typical resistance of ∼2 M in the bath solution. The bath solution contained 130 mM NaCl, 5 mM KCl, 2 mM CaCl_2_, 1 mM MgCl_2_, 10 mM HEPES-NaOH, and 10 mM glucose; the pH was adjusted to 7.3 with NaOH. The solution in the cell-attached pipette contained 40 mM NaCl, 100 mM TEACl, 5 mM KCl, 2 mM CaCl_2_, 1 mM MgCl_2_, and 10 mM HEPES-NaOH; the pH was adjusted to 7.3 with NaOH. As mentioned previously (Yao et al., 2013; Yao et al., 2012), capacitance steps and fission-pore durations were considered as reliably detected for step sizes >0.2 fF, and smaller capacitance steps are not included in the analysis. The number of endocytic events per patch detected in the cell-attached recordings is counted as the total number of downward capacitance steps within the first 5 min of recordings (Yao et al., 2013; Yao et al., 2012).

The fusion-pore conductance (Gp) for exocytic events or the fission-pore Gp for endocytic events can be calculated from both Re and Im traces as 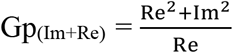 or from Im trace alone as 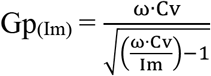, as described (Gong et al., 2007; Yao et al., 2013; Yao et al., 2012). Analysis of fission-pore kinetics were restricted to fission-pores with durations >15 ms, since shorter events were distorted by the lock-in amplifier low-pass filter (set to 1 ms, 24 dB) (Yao et al., 2013; Yao et al., 2012).

### Double (whole-cell/cell-attached) patch recordings in chromaffin cells

Double (cell-attached/whole-cell) patch recordings were performed to analyze endocytosis-associated currents, in which the cell-attached patch pipette was utilized to simultaneously monitor endocytic events and the ionic current across the patch membrane. With the cell constantly clamped at −60 mV by the whole-cell patch pipette via an EPC-10 amplifier (HEKA Elektronik, Germany), the patch membrane was held at different voltages by varying the voltage in the cell-attached pipette. The solution in the whole-cell pipette contained 20 mM CsCl, 90 mM KCl, 20 TEACl, 10 mM HEPES-KOH, 2 mM EGTA, 2 mM Na_2_ATP, and 0.3 Na_3_GTP; the pH was adjusted to 7.3 with KOH with the osmolarity of 290 mmol kg^−1^. The basal Ca^2+^ level in the whole-cell pipette solution was buffered by 1.1 mM CaCl_2_ and 2 mM EGTA to give a final free Ca^2+^ level of ∼150 nM. All the settings for cell-attached recordings and the solution in the cell-attached pipette were identical to descriptions in “*Cell-attached capacitance recordings and analyses of exocytic fusion-pore and endocytic fission-pore*”. Ionic currents from the cell-attached pipette were low-pass filtered at 500 Hz. As illustrated in Fig. 1F, the endocytosis-associated current was calculated as the difference between linear fittings of a 20-100 ms duration in the I_patch_ trace before and after the capacitance drop due to endocytosis.

### Paired whole-cell recordings of evoked IPSCs in neurons

Paired whole-cell patch-clamp recordings were performed on DIV 14-16 neurons, continuously perfused with an extracellular solution containing 130 mM NaCl, 3 mM KCl, 10 mM HEPES-NaOH, 2 mM CaCl_2_, 1 mM MgCl_2_ and 10 mM glucose, pH adjusted to 7.3 with NaOH. Evoked IPSCs were isolated by adding amino-5-phosphonopentanoate (AP5, Tocris) at 50 μM and 6-cyano-7-nitroquinoxaline-2,3-dione (CNQX, Tocris) at 20 μM in the bath solution to block excitatory synaptic transmission. The pipette solution for presynaptic neurons contained 125 mM KCl, 10 mM HEPES, 2mM EGTA-KOH, 1mM CaCl_2_, 1mM MgCl_2_, 2 mM Na_2_ATP, 0.4 mM Na_3_GTP and 5 mM phosphocreatine disodium salt, the pH adjusted to 7.3 with KOH. For postsynaptic neurons, the pipette solution contained 125 mM CsCl, 10 mM HEPES, 2 mM EGTA-CsOH, 1 mM CaCl_2_, 1 mM MgCl_2_, 2 mM Na_2_ATP, 0.4 mM Na_3_GTP, 5 mM QX-314, and 5 mM phosphocreatine disodium salt, pH adjusted to 7.3 with CsOH.

IPSCs were recorded at −70 mV on pyramidal neurons using dual whole-cell configurations by evoking a nearby inhibitory neuron with a 1-ms depolarization from −70 to +30 mV. Recordings were acquired with an EPC-10 patch-clamp amplifier for the presynaptic voltage clamping and an EPC-7 plus patch-clamp amplifier for postsynaptic recordings. Data were filtered at 1 kHz and digitized at 5 kHz. Patch pipettes had a typical resistance of 2-3 MΩ. Criteria for inclusion required that Rs remained below 10 MΩ and did not increase by more than 20% during the course of recordings.

Analysis was carried out with Igor software (Wavemetrics), using custom-written analysis procedures. Peak amplitudes of inhibitory postsynaptic currents (IPSCs) were extracted by subtracting the extrapolated residual current caused by previous stimulation. The amplitude of IPSCs during train stimulations was normalized by dividing the peak of each response to the peak amplitude of the first response.

### Whole-cell recordings of TRPM7 currents in HEK293 cells

TRPM7 currents were monitored in HEK293 cells using the whole-cell configuration as previous reported (Aarts et al., 2003). The bath solution contained 130 mM NaCl, 5 mM KCl, 1 mM CaCl_2_, 1 mM MgCl_2_, 10 mM HEPES-NaOH, and 10 mM glucose, 0.1 mM CdCl_2_, 1 µM TTX; the pH was adjusted to 7.3 with NaOH, and the osmolarity was ∼310 mmol kg^−1^. The solution in the whole-cell pipette contained 120 mM Cs-MeSO_3_, 8 mM NaCl, 2 mM CaCl_2_, 10 mM EGTA-CsOH and 10 mM HEPES; the pH was adjusted to 7.3 with CsOH, and the osmolarity was adjusted to 290 mmol kg^−1^. The voltage ramp protocol holds cells at 0 mV, steps to −100 mV for 40 ms and then ramps to +100 mV over 500 ms, holding at +100 mV for 40 ms before stepping back to 0 mV, and this protocol is repeated every 5 s during recording. Series resistance was monitored by applying a voltage step of 10 mV. Current signals were filtered at 3 kHz and digitized at 5 kHz. Only recordings with a pipette-membrane seal resistance >2GΩ were included. Current traces and time courses were analyzed with customized macro for Igor software (WaveMetrics, Lake Oswego, OR).

### Carbon fiber amperometry

Chromaffin cells were incubated in the bath solution containing 140 mM NaCl, 5 mM KCl, 2 mM CaCl_2_, 1 mM MgCl_2_, 10 mM HEPES-NaOH, and 10 mM glucose, pH 7.3 with NaOH. The stimulation solution contained 45 mM NaCl, 90 mM KCl, 2 mM CaCl_2_, 1 mM MgCl_2_, 10 mM HEPES-NaOH, and 10 mM glucose. Conventional carbon fiber amperometry for catecholamine detection used 5-µm carbon fibers (ALA Scientific, Farmingdale, NY) as described previously (Gong et al., 2007; Gong et al., 2005; Yao et al., 2012). The freshly cut tip of the carbon fiber electrode was positioned closely against the cell surface to minimize the diffusion distance from release sites, and cells were stimulated with 90 mM KCl solution for 5 s by a pressurized perfusion system (VC3, ALA scientific) placed ∼40 µm away from the cell. The amperometric current, generated by catecholamine oxidation at the exposed tip of the carbon fiber electrode due to stimulation, was measured using an EPC-7 plus amplifier at a holding potential of +700 mV. Amperometric signals were low-pass filtered at 3 kHz and digitized at 5 kHz. Amperometric recordings were collected and then analyzed with a customized macro for Igor software (WaveMetrics, Lake Oswego, OR) to extract spike information according to the criteria of Chow et al (Chow et al., 1992). The following criteria were set in single-spike analysis: (i) only spikes >10 pA were considered; (ii) the maximum number of spikes analyzed per cell was set to 100; (iii) foot duration was delimited by the baseline and spike onset as shown previously (Chow et al., 1992), and (iv) spikes with a foot duration of < 0.5 ms were excluded for the assays. Data are presented as mean ± SEM of the cell medians. Thus, the numbers used for statistical tests are the number of cells. The number of amperometric spikes was counted as the total number of spikes with an amplitude >10 pA within 15 s after stimulation.

### Live-cell imaging

Neurons on coverslips were continuously perfused at a flow rate of ∼1 ml/min with bath solution (130 mM NaCl, 2.8 mM KCl, 2 mM CaCl_2_, 1 mM MgCl_2_, 10 mM glucose, 10 mM HEPES [pH 7.4]; ∼310 mOsm). All imaging experiments were performed on an Olympus IX51 microscope with a 60x oil immersion objective at room temperature (25 ± 1 °C), with a GFP-3035B filter cube (472/30 nm excitation, 520/35 nm emission, 495 nm dichroic mirror) (Semrock, Rochester, NY) as the filter set. To elicit responses, action potentials (APs) were delivered by electric field stimulation at 10V/cm with 1-ms duration via two parallel platinum wires embedded within the perfusion chamber with 7-mm spacing. 20 μM 6-cyano-7-nitroquinoxaline-2,3-dione (CNQX, Tocris Bioscience) and 50 μM D, L-2-amino-5-phosphonovaleric acid (AP5, Tocris Bioscience) were included in the bath solution to prevent recurrent activities.

For train stimulation experiments using sypHy, vGlut1-pHluorin, or SyGCaMP6f, a 120-W mercury vapor short Arc lamp (X-cite, 120PC, Exfo) was utilized as the light source for excitation. Images were captured by a Photometrics HQ2 coolSNAP CCD camera controlled by μManager software (micro-manager.org) at a 1-2 s interval with 100 or 200-ms exposure time at 25% illumination. To measure reacidification rate of newly formed endocytic vesicles using sypHy as described previously (Atluri and Ryan, 2006), neurons on coverslip were rapidly superfused with a valve controlled pressurized perfusion system (ALA Scientific, Farmingdale, NY). The perfusing buffer was switched between the standard pH 7.3 bath solution described above and pH 5.5 buffer with the HEPES substituted with equi-molar MES. The electrical stimulation was thus time-locked to frame acquisition and buffer changes.

For single AP stimulation using SyGCaMP6f and train stimulation using iGluSnFR, Lambda 421 Optical Beam Combining System (Sutter instrument, Novato, CA) was used as the light source for illumination. To achieve high quantum efficiency, we used the Prime BSI Scientific CMOS (sCMOS) camera (Teledyne Photometrics). With 20-ms exposure time, the illumination via Lambda 421 and data acquisition via camera were controlled by Igor software (WaveMetrics, Lake Oswego, OR) to achieve a fast sampling rate of 50 Hz. The light intensity of illumination was set at 10% to minimize photobleaching.

### Image analysis

Images were analyzed in ImageJ <u>(ht</u>t<u>p://rsb.info.nih.gov/ij)</u> using custom-written plugins <u>(ht</u>t<u>p://rsb.info.nih.gov/ij/plugins/time-series.html)</u> and Igor software (Wavemetrics, Lake Oswego, OR) using custom-written procedures. Photobleaching was less than 2% for all images included within analysis and thus was not corrected. To analyze images from sypHy, vGlut1-pHluorin, or SyGCaMP6f experiments, regions of interest (ROIs) of identical size (4 x 4 pixels) were placed in the center of individual synaptic boutons reacting to stimuli, and fluorescence changes were tracked throughout the image stack. In general, 10-20 ROIs were analyzed from each coverslip for sypHy or vGlut1-pHluorin experiments, and 20-40 ROIs for all SyGCaMP6f experiments. For iGluSnFR analysis, ROIs were created as previously reported (Vevea and Chapman, 2020). Briefly, ROIs (>10 pixels), defined by a series of image subtractions and thresholding (Vevea and Chapman, 2020), were used to measure fluorescence changes of image stacks.

Fluorescent intensities of individual ROIs prior to stimuli were averaged as baseline (F0). The fluorescent changes (ΔF) were normalized to the peak of fluorescent increase (ΔFmax) as ΔF/ΔFmax for sypHy, vGlut1-pHluorin or iGluSnFR experiments and as ΔF/F0 for SyGCaMP6f experiments. Normalized ΔF of individual ROIs were taken as independent replicates (n) for sypHy and vGlut1-pHluorin experiments with the average of normalized ΔF of ROIs for each coverslip as the independent replicates (n) for SyGCaMP6f and iGluSnFR experiments.

### DNA constructs

GFP in the lentiviral vector (pCDH-EF1-MCS-T2A-GFP) was removed and replaced with an artificial DNA fragment containing *Pme* I and *Apa* I restriction sites using sense oligo: 5’-CCGGAATGTTTAAACGGGCCCG and antisense oligo: 5’-TCGACGGGCCCGTTTAAACATT-3’. To engineer pCDH-EF1-sypHy, sypHy (#24478 of Addgene, a kind gift from Dr. L. Lagnado (Granseth et al., 2006)) was inserted into MCS by adding an *Xba* I site to the 5’ end (primer: 5’-CTTTTCTAGAATGGACGTGGTGAATCA-3’) and a *Not* I site to the 3’ end (primer: 5’-CAGCGGCCGCCATCTGATTGGAGAAGGAG-3’). The EF1 promoter was then replaced with a human synapsin 1 (SYN1) promoter to produce pCDH-SYN1-sypHy, which was packaged into lentivirus and used for neuronal infections.

SyGCaMP6, a kind gift from Dr. Timothy A. Ryan’s lab, was generated by adding GCaMP6f to the C terminus of the mouse sequence of synaptophysin as previously reported (Kim and Ryan, 2013). SyGCaMP6f was first cloned by the standard PCR protocol using Phusion DNA polymerase (forward primer: 5’-ACTCGGATCCCCTCTAGAATGGACGTGGTGAATCAGCTG-3’; reverse primer: 5’-GATTGATATCTCACTTCGCTGTCATCATTTG-3’). The PCR fragment was then enzyme digested and ligated between the *BamHI* and *EcoRV* sites to obtain pCDH-SYN1-SyGCaMP6f by replacing sypHy in the pCDH-SYN1-sypHy vector.

Modified iGluSnFR (Marvin et al., 2018), with the final four amino acid removed and a Golgi export sequence and ER exit motif included, is a kind gift from Dr. Edwin Chapman’s lab (Bradberry et al., 2020). iGluSnFR was subcloned and inserted between the BamH1 and Not1 restriction sites in the lentiviral vector (pCDH-CaMKIIα -MCS, a construct we created in lab) to obtain pCDH-CaMKIIα-iGluSnFR (forward primer: 5’-ATAAGGATCCATGGAGACAGACACACTCCTGCTATG-3’, and reverse primer: 5’-CGATGCGGCCGCTTAGAGGGCAACTTCATTTTCATAGC −3’).

To express TRPM7 in KO chromaffin cells with lentivirus, TRPM7 (#45482 of Addgene, a kind gift of Dr. A. Scharenberg’s lab (Nadler et al., 2001)) was subcloned into a lentiviral vector via *Pme* I and *Apa* I restriction sites to generate pCDH-EF1-TRPM7^WT^.

To engineer TRPM7 mutant with a loss of channel function (TRPM7^LCF^, amino acid 1090-1092, NLL>FAP) (Krapivinsky et al., 2006), site-directed mutagenesis was performed using *Pfu* hotstart DNA polymerase (forward primer: 5’-CAGTATATCATTATGGTTTTCGCTCCTATCGCATTTTTCAATAAT-3’, and reverse primer: 5’-ATTATTGAAAAATGCGATAGGAGCGAAAACCATAAT GATATACTG-3’). Meanwhile, to express TRPM7 in KO neurons, TRPM7^WT^ or TRPM7^LCF^ was digested out with *Pme* I and *Apa* I restriction sites and ligated into pCDH-EF1-sypHy with the artificial DNA fragment to produce pCDH-EF1-sypHy-T2A-TRPM7.

### Lentiviral productions

Low-passage HEK293 cells (Thermo Fisher) were co-transfected with 10 μg lentiviral vector (pCDH-SYN1-sypHy, pCDH-SYN1-SyGCaMP6f, pCDH-CaMKIIα-iGluSnFR, pCDH-EF1-TRPM7^WT^ or pCDH-EF1-TRPM7^LCF^) along with 7 μg packaging vector psPAX2 and 3 μg envelope vector pMD2.VSVG using the polyethylenimine (PEI) mediated transfection method. The supernatant containing lentiviral particles was collected and filtered through 0.45 μm filter to remove cell debris at 48 and 72 hrs. Lentiviral particles, concentrated by PEG-it™ precipitation kits from System Bioscience (Palo Alto, CA), were re-suspended in cold PBS and stored at −80 °C.

There is a logarithmic inverse relationship between packaging size and lentiviral titer, that is, the titer drops by 1 log for every 2kb of insert (Kumar et al., 2001). Since *Trpm7* is a relatively large gene of >5.5 kb, to increase the packaging efficiency, *Flag* was tagged to the 5’-end of *Trpm7* variants for lentivirus, which was utilized to infect KO chromaffin cells. Our experiments showed that the majority of chromaffin cells were infected by anti-FLAG immunostainings (94 ± 1.4% from 3 independent experiments).

### Statistical analysis

Data were tested for normal distribution and, if necessary, log-transformed for the fission-pore duration in Fig. 1E and 4G to fulfill the criteria of normal distributions for student’s t-tests. All analyses were performed with SPSS.

No statistical method was used to predetermine sample size, but our samples sizes are similar to those reported in previous studies (Fernandez-Alfonso and Ryan, 2004; Granseth et al., 2006; Soykan et al., 2017; Yao et al., 2013; Yao et al., 2012). Data was expressed as mean ± s.e.m, animals or cells were randomly assigned into control or experimental groups. Statistical analysis was performed with unpaired two-tailed student’s t-test except paired two-tailed student’s t-test in Supplemental fig. 2B and Newman of one-way ANOVA in Fig. 5B and Supplementary fig. 4B.

## Acknowledgements

This work is supported by NIH (R01NS110533) to L.-W.G. and NSF (IOS-1346826) to B.S.G.. The authors are grateful to Dr. V. Haucke for vGlut1-pHluorin, Dr. T.A. Ryan for SyGCaMP6f, Dr. E.R. Chapman for iGluSnFR, and Dr. A.M. Scharenberg for TRPM7 plasmid (#45482, Addgene).

## Author contributions

Z.-J.J., W.L., L.-H.Y. and L.-W.G. conceived the idea. Z.-J.J., L.-H.Y. and K.V. carried out electrophysiological experiments. Z.-J.J., W.L., A.M. and S.A. performed live-cell imaging experiments. B.S.G. carried out RT-PCR experiments. Z.-J.J. and L.-W.G. wrote the paper. All authors contributed to the discussion and revision of the paper.

## Resource sharing

Further information and reasonable requests for resources and reagents should be directed to the lead contact, Liang-Wei Gong.

## Supplementary information

**Supplementary fig 1.**
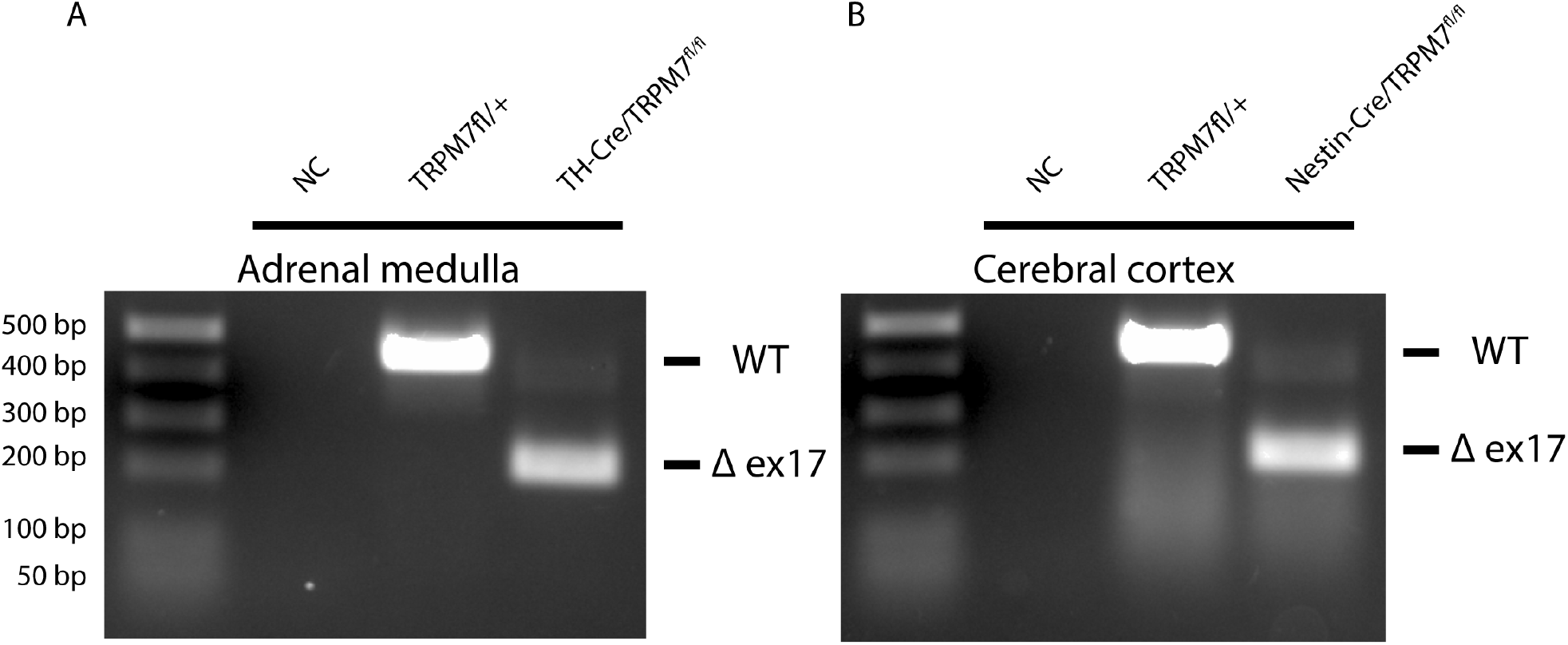
TRPM7 depletion in cortex of the brain and adrenal medulla. (**a-b**) RT-PCR of total RNA from adrenal medulla (A) and cortex (B) showing presence or absence of exon 17 of *trpm7* in the differentially-sized amplicons.

**Supplementary fig 2.**
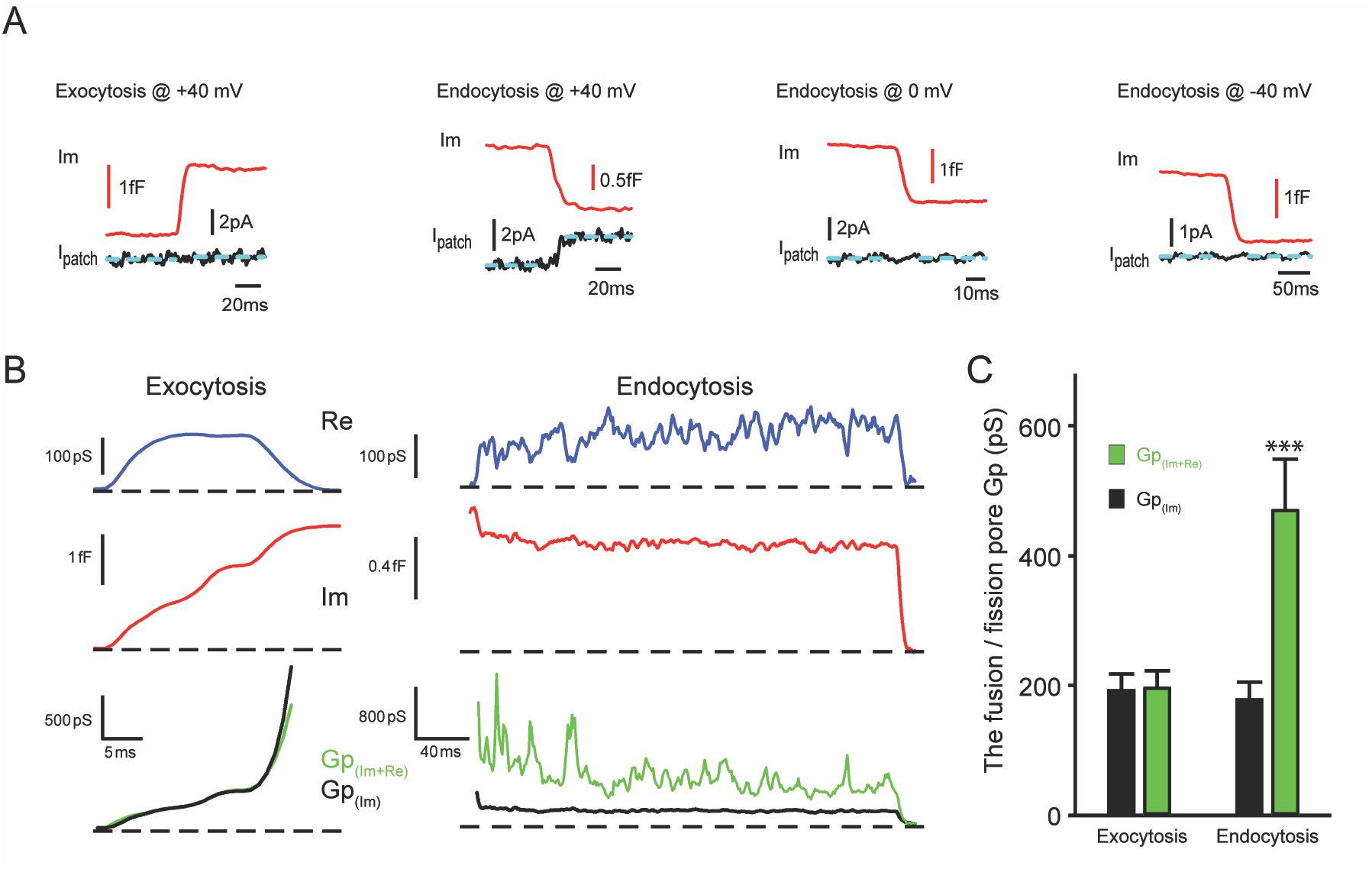
An ionic current is specifically associated with single vesicle endocytosis. (**A**) Representative membrane capacitance (Im) and patch membrane current (I_patch_) traces for exocytosis or endocytosis at the indicated membrane potentials. Dashed lines in cyan represent fitting line before or after capacitance increase for exocytosis or decrease for endocytosis. **(B)** Representative exocytic (left) and endocytic (right) events shown as (from top down) membrane conductance (Re), capacitance (Im), and pore conductance (Gp). Gp_(Im+Re)_ (Green) is calculated from both membrane conductance and capacitance traces and Gp_(Im)_ (Black) is calculated from the capacitance trace, as indicated in methods. Dashed lines represent the baselines. **(C)** The statistical analysis of Gp_(Im+Re)_ and Gp_(Im)_ for exocytosis and endocytosis indicates the presence of an extra conductance and thus an ionic current associated with endocytosis but not exocytosis (Exo: p = 0.99, n = 15; Endo: p < 0.001, n = 13).

**Supplementary fig 3.**
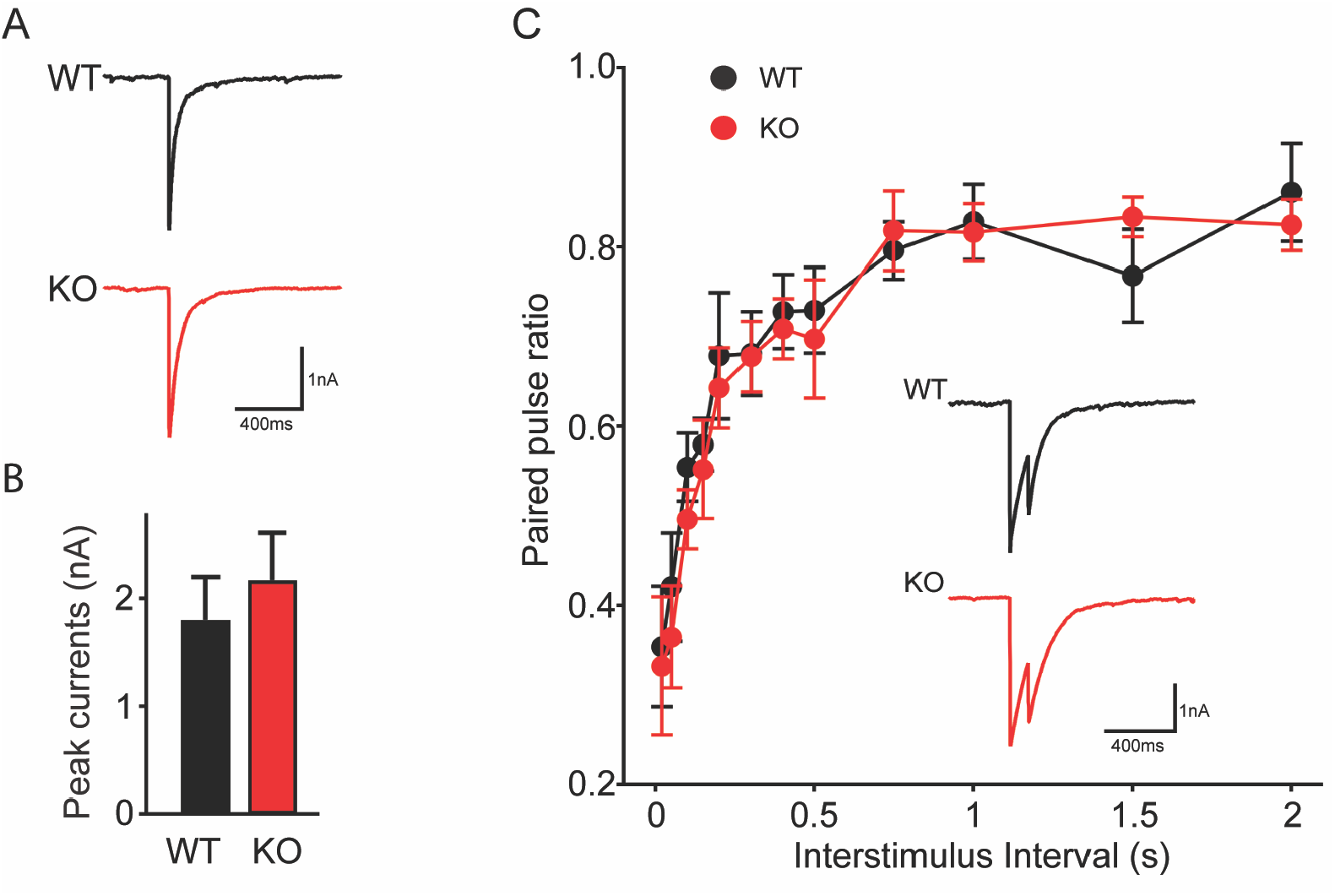
TRPM7 may not be involved in synaptic vesicle exocytosis in inhibitory neurons. (**A-B**) Representative (a) and quantification (b) of IPSCs evoked by a single stimulation in WT and TRPM7 KO neurons (WT: n = 14; KO: n = 13). (**C**) Quantification of the paired pulse ratio with variable intervals between WT and KO neurons (WT: n = 12; KO: n = 12), and insert is representative IPSCs evoked by paired stimuli at 100-ms intervals.

**Supplementary fig 4.**
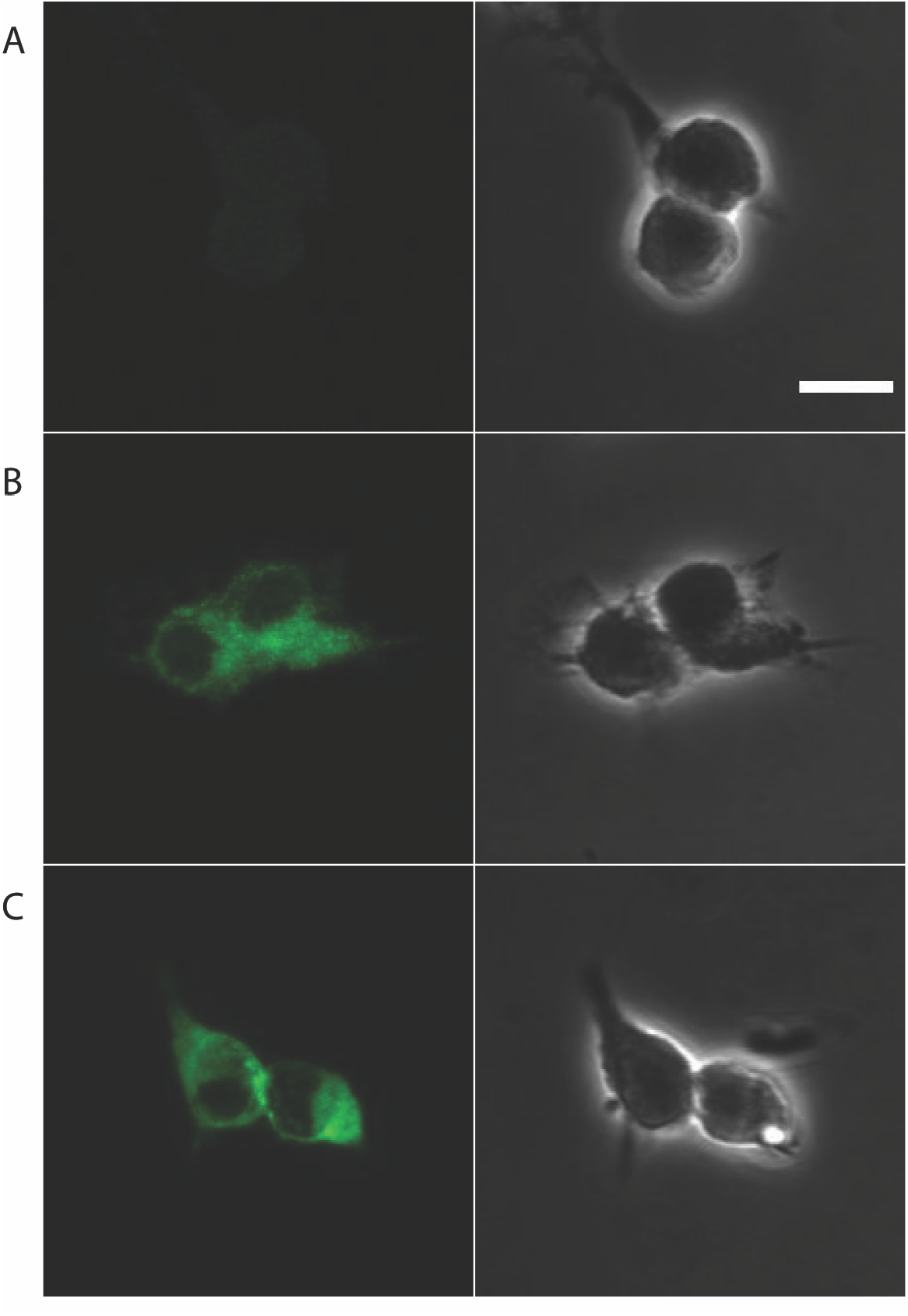
Confocal images for HEK293 cells infected with lentivirus encoding with either empty vector, TRPM7^WT^ or TRPM7^LCF^. (**A-C**) Representative confocal images of HEK293 cells infected with lentivirus encoding empty vector (A), TRPM7^WT^ (B) or TRPM7^LCF^ (C) are on the left, phase contrast images of the same ROIs are on the right. The scale bar is 10 μm.

**Supplementary fig 5.**
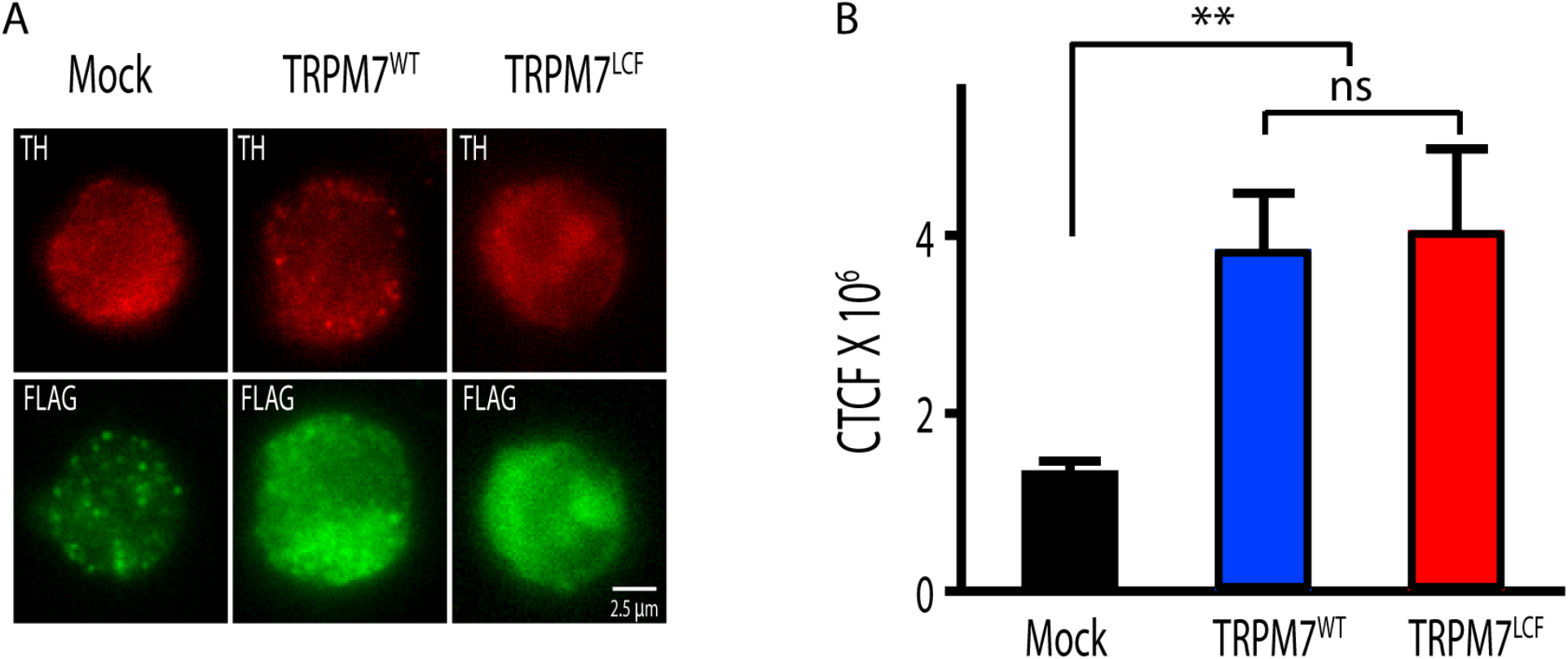
Similar expression levels of TRPM7^WT^ and TRPM7^LCF^ in KO chromaffin cells. (**A**) Representative images of TRPM7 KO cells, which were infected with lentivirus encoding empty vector (left), pCDH-EF1-flag-TRPM7^WT^ (middle) or pCDH-EF1-flag-TRPM7^LCF^ (right), immunostained for tyrosine hydroxylase (TH) (top) and FLAG (bottom). Bar = 2.5 μm. (**B**) Quantification of whole cell anti-FLAG immunofluorescence in these 3 groups indicates similar expression levels of TRPM7^WT^ and TRPM7^LCF^ in KO cells. Fluorescence intensity was analyzed as CTCF (the corrected total cell fluorescence), which was calculated using the formula: CTCF = integrated density − (area of selected cell × mean fluorescence of background readings). n =10 cells for each group, and a.u. stands for arbitrary units. * * p < 0.01 and N.S., non-significant, Newman of one-way ANOVA.

